# G-S-M: A Comprehensive Framework for Integrative Feature Selection in Omics Data Analysis and Beyond

**DOI:** 10.1101/2024.03.30.585514

**Authors:** Malik Yousef, Jens Allmer, Yasin İnal, Burcu Bakir Gungor

**Affiliations:** Department of Information Systems, Zefat Academic College, Zefat, Israel; Medical Informatics and Bioinformatics, Institute for Measurement Engineering and Sensor Technology, Hochschule Ruhr West, University of Applied Sciences, Mülheim an der Ruhr, Germany; Department of Computer Engineering, Abdullah Gul University, Kayseri, Türkiye

## Abstract

The treatment of human diseases is a major research question in many fields related to medicine. It has become clear that patient stratification is of utmost importance so that patients receive the best possible treatment. Bio/disease markers are critical to achieve stratification. Markers can come from many different sources such as genomics, transcriptomics, and proteomics. Establishing markers from such measurements often involves data analysis, machine learning, and feature selection. Traditional feature selection techniques often rely on the estimation of individual feature importance or significance by assigning a score to each feature, disregarding the inter-feature relationships. In contrast, the G-S-M (grouping scoring modeling) approach considers a group of features as a set that is organized based on prior knowledge. This approach takes into account the interdependence among features, providing a more meaningful evaluation of feature relevance and utility. Prior knowledge can encompass much compiled information such as microRNA-target interactions and protein-protein interactions. Here we present a new tool called G-S-M that presents the generalization of our previous works such as maTE, CogNet, and PriPath. The G-S-M tool combines machine learning and prior knowledge to group and score features based on their association with a binary-labeled target such as control and disease. This approach is unique in that computational and domain knowledge is utilized concurrently. Embedded feature selection, repeatedly employing machine learning during the selection process results in the identification of the most discriminative groups.

Furthermore, the G-S-M tool allows for a more holistic understanding of the underlying mechanisms of a given system to be achieved through the combination of machine learning and prior domain knowledge, which can lead to new insights and discoveries. The implementation of the G-S-M workflow is freely available for download from our GitHub repository: https://github.com/malikyousef/The-G-S-M-Grouping-Scoring-Modeling-Approach. With this generalized approach we aim to make the feature selection approach available to a broader audience and hope it will be employed in medical practice. An example of such an approach is the TextNetTopics that is based on the G-S-M approach. TextNetTopics uses Latent Dirichlet Allocation (LDA) to detect topics of words, where those topics serve as groups. In the future, we aim to extend the approach to enable the incorporation of multiple lines of evidence for biomarker detection and patient stratification via combining multi-omics data.

## Introduction

The complex relationship among environment, genes, and gene expression reveals itself in disease - drug - molecular interactions that are important in precision medicine. Based on the genetic make-up and the current expression levels of genes (RNA and protein) some drugs may be more effective than others for the same phenotype or may be very detrimental. Such relationships can be investigated on the molecular level using various omic technologies such as transcriptomics, proteomics, and lipidomics. Employing the knowledge at the bedside requires clinically specific bio- and disease markers that can delineate among phenotypes and how to treat them.

Hence, the identification of biomarkers for various diseases has been a subject of great interest within the field of bioinformatics and machine learning. Much information has been accrued and is stored in various databases such as DisGeNET, OMIM, and RNAcentral. For a review of how RNA databases can be manually utilized in cancer research, have a look at Allmer 2023. A significant challenge in bioinformatics is to incorporate such information in a consistent, automated, and meaningful manner. Several approaches have been developed, such as statistical, network-based, and machine learning-based methods. In bioinformatics, tools like gene ontology enrichment analysis, DAVID^1^, STRING^2^, and Cytoscape^3^ have been employed to analyze complex biological data.

Another method for integrating previous knowledge using various computational tools is the grouping, scoring, and modeling (G-S-M) approach. This approach combines molecular measurements with prior knowledge, such as miRNA regulation, KEGG pathways, and disease databases. This incorporation of prior knowledge can be accomplished via embedded feature selection with machine learning at the heart of the algorithms. Implementations of the G-S-M approach such as CogNet^4^, maTE^5^, mirCorrnet^6^, miRModuleNet^7^, SVM-RCE-R^8^, PriPath^9^, and GediNET^10^ served as the inspiration for the current study. The G-S-M algorithm considers groups of gene expressions rather than assuming an individual impact of gene expression on a phenotype. For each group G-S-M extracts the sub_dataset from the original dataset and then applies an internal cross-validation to assign a score for the group. During this process, it also selects features based on the numeric values by performing a t-test between classes to make the process more discriminative. Classes in machine learning are annotations that are attached to the samples and that differentiate among samples so that a model can be trained to differentiate among samples given the class label (its annotation). Typically labels are positive and negative in machine learning, but they can be anything else such as target and control so we use the term annotation instead of the term class label to indicate this freedom.

The primary objective of this study is to build on top of our previous works and abstract from any particular prior knowledge. In our previous applications we considered specific prior knowledge such as miRNA-target interactions in maTE, disease genes associates in GediNET^11^, and KEGG pathway in PriPath^12^. Here we present a generalization of the G-S-M approach which can be applied to any form of prior knowledge that can group the measured features. To achieve this goal, we implemented a new KNIME workflow entitled G-S-M. It utilizes two-class classification and, therefore, needs data from two classes e.g., control and disease. Additionally, the prior knowledge, e.g., genes associated with a disease, is needed as input. Grouping is then performed according to the prior knowledge. However, the t-test is applied on the training set to reduce the number of features and also provide significant features to the Grouping component.

Embedded feature selection for training testing is then performed using Monte Carlo cross validation (MCCV). In each training iteration, the groups with the most accurate results are determined and the highest-ranked groups are used to train the model. Over multiple rounds of MCCV, the models are ranked on their average performance. The groups are then ranked by their model performance. This effectively selects for the groups, that group features by prior knowledge, that best discriminate between the classes. This informed embedded feature selection approach has been shown to work well in practice (e.g., maTE and PriPath). For the G-S-M generalization, we reproduced the results for some of our previous works. We were able to largely reproduce the previous results and conclude that the generalization of the G-S-M approach to arbitrary prior knowledge has been successfully achieved.

In the future, we aim to extend this approach to accept multiple measurements from different omics levels. Thereby, we hope to improve patient stratification and to abstract from the particular possibilities of the specific clinic that performs it, making the tool universally applicable from using regular blood work to including multi-omics information, including transcriptomic, proteomic, and metabolomic data. Together, this will make patient stratification more accessible in practice.

## Materials and Methods

### Necessary Input for the G-S-M tool

For the tool to function effectively, it requires two distinct files: a primary input data file, and a secondary grouping file that contains pre-existing knowledge.

The input data file is the data provided by the user, typically in the form of a tabular structure. The table (Table X) should contain the attributes or features that are intended to be used as columns, as well as a designated target column that is comprised of two distinct class labels. Additionally, it is important to note that the file extension for this input data file must be “.csv” for it to be compatible with our KNIME^13^ workflow for processing.

**Table X:** An abridged version of the sample_measurements.csv file. Each symbol can only have one measurement in the file. The file needs to be set in the workflow or needs to be placed with the exact name in the root of the WF directory.

|  | Annotation | symbol 1 | symbol 2 | symbol n |
| --- | --- | --- | --- | --- |
| sample 1 | ..... | ..... | ..... | ..... |
| sample 2 | ..... | ..... | ..... | ..... |
| sample n | ..... | ..... | ..... | ..... |

The second file that is required for the grouping process is a file that contains pre-existing information. The structure of this file should consist of group names and their corresponding desired attributes (Table Y).

**Table Y:** An abridged version of the prior-info.csv file that needs to be provided to the G-S-M workflow. The relationship between symbol and group is many to many. The file needs to be set in the workflow or needs to be placed with the exact name in the root of the WF directory. The example contains gene symbols (Symbol) and diseases that group the genes (Group).

| Symbol | Group |
| --- | --- |
| PLXNA3 | Endometrial Carcinoma |
| PLXNA3 | Breast Carcinoma |
| PLXNA3 | Liver carcinoma |
| CDC42BPG | Glioblastoma Multiforme |
| DNAH3 | Colorectal Carcinoma |
| GALNT10 | Metastatic Renal Cell Cancer |
| GALNT10 | Liver carcinoma |
| GALNT10 | Metastatic Renal Cell Carcinoma |
| CDV3 | Colorectal Carcinoma |

### Datasets for G-S-M Tests

There are two kinds of resources used by G-S-M, the prior-info file and the expression measurements. The prior-info derived from a,b,c,.. to represent diseases, miRNAs.The gene expression datasets were downloaded from GEO. We have used the GDS196 data to test the workflow. Other datasets were used in related studies that use the G-S-M approaches. Table Z demonstrates different studies that utilized the G-S-M approach with different prior knowledge for grouping..

**Table Z:** It demonstrates different studies that utilized the G-S-M approach with different grouping factors.

| Project Title | Prior Knowledge | Example Group | Reference |
| --- | --- | --- | --- |
| GediNET- Discovering Multi-Disease Gene Associations | Disease associations | Malignant Glioma = {TK1, NPAS3, CD63, HMGB1..} | 11 |
| PriPath: Identifying Dysregulated Pathways from Differential Gene Expression via Grouping, Scoring, and Modeling with an Embedded Feature Selection Approach | KEGG Pathways | Chronic myeloid leukemia = { TP53, MYC, HRAS, KRAS...} | 12 |
| miRDisNet: A novel approach for cancer classification using mir-disease associations | miRNA - disease | Acute Brucellosis = { hsa-mir-126, hsa-mir-4753 } | 15 |
| TextNetTopics: Text Classification Based Word Grouping as Topics and Topics' Scoring | LDA Topics | topic_0 = "patient, disease, studi, risk, rate, percent, transplant, clinic, treatment, compar" | 16 |

### The G-S-M Approach

The G-S-M approach employed by G-S-M Tool is a generalization of the previous implementation in our tools such as maTE and GediNET. **Figure 2** illustrates the basic flow of the G-S-M Tool process, with the green section representing the Grouping component, the orange section representing the Scoring component, and the blue section representing the Modeling component.

**Figure 2.**
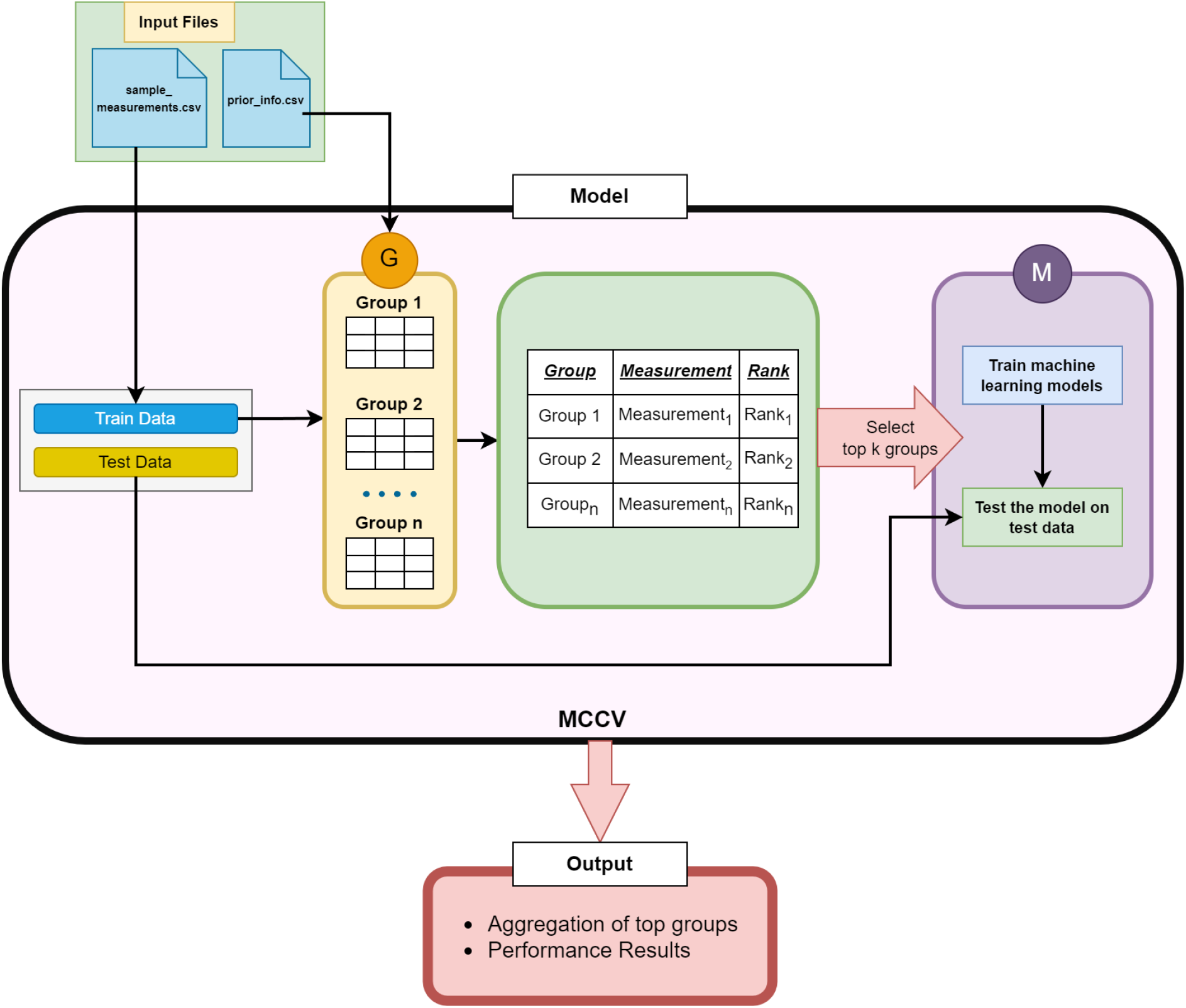
shows the workflow of the G-S-M Tool. The main process of G-S-M integrates previous biological information to classify genes based on their relationship to a grouping factor (e.g., diseases). This data is provided by the user.

**Figure 3:**
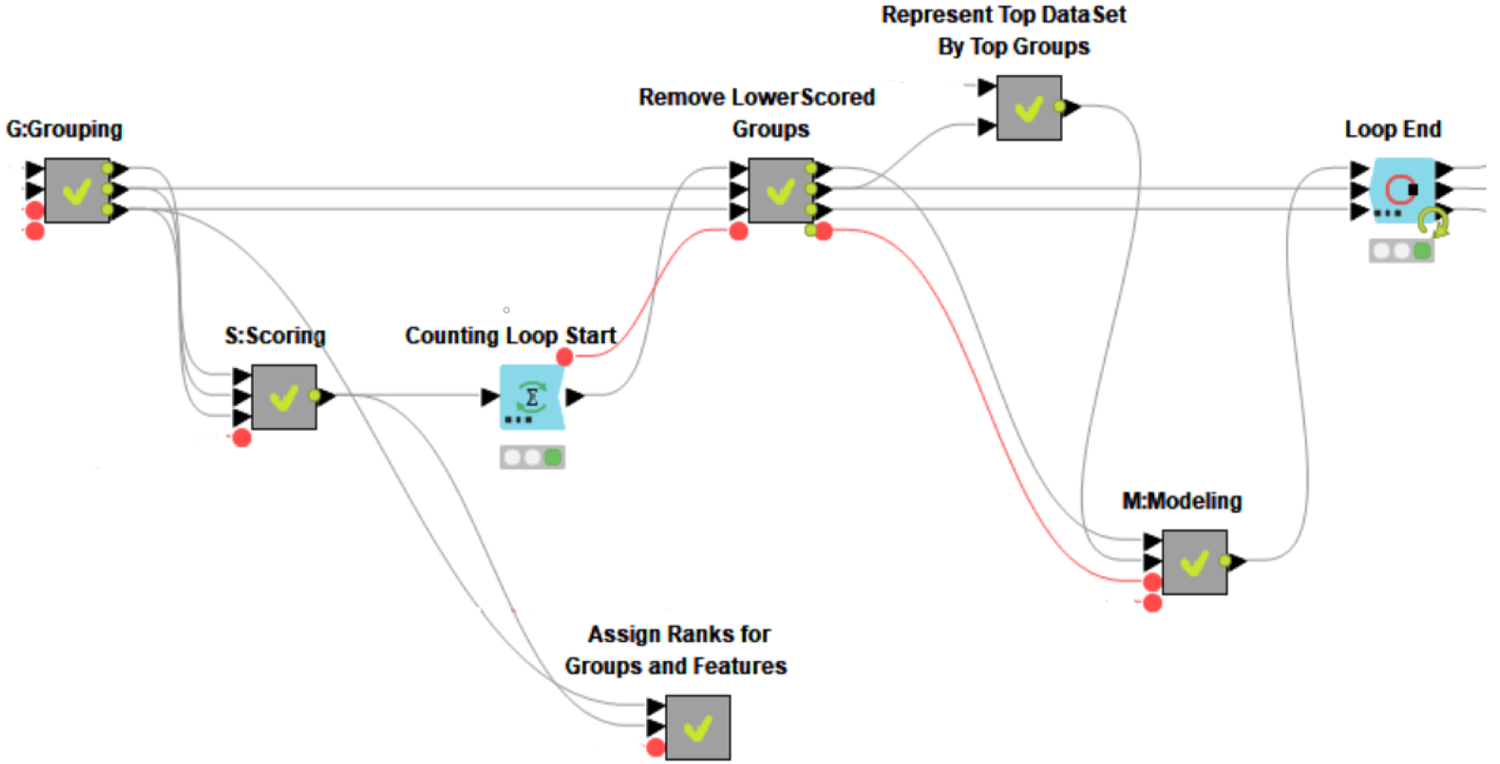
The core of the GSM workflow. G, S, and M are indicated. The data processing is performed in the meta-nodes which can be expanded to inspect the workflow further. The user does not have to interact with the workflow on this level but can set the necessary input files and parameters on a higher level.

1. The G Component (Grouping): responsible for organizing the attributes in the main dataset into smaller groups based on pre-existing knowledge about their potential role in disease. Each group is represented by a subset of the data that only includes two categories or classes, and these sub-datasets are extracted from the main dataset.
2. The S Component (Scoring): responsible for evaluating and ranking the groups of features based on their corresponding two-class sub-datasets. The scores are calculated by considering the information and characteristics of these sub-datasets.
3. The M Component (Modeling): responsible for creating a model by training a classifier (such as Random Forest, Decision Tree, Naive Bayes, Support Vector Machine, or Random Forest-based TreeBag) on the predictors from the highest-ranked groups. The classifier is used to make predictions about the relationships between the predictors and the target.

G-S-M Tool is a tool that utilizes a two-class gene expression dataset and a table containing existing knowledge about the diseases being studied. The samples in the dataset are divided into two categories: control (negative, healthy) and disease (positive, patient). The dataset is split into training and testing sets, with the training set being used for the G-S-M components and the testing set being used to evaluate the performance of the model. This process is repeated n (user choice) times using cross-validation, where the input is randomly divided into 90% training and 10% testing in each iteration. A statistical t-test (one-way ANOVA) is applied to the training dataset to identify the most differentially expressed genes. The top n genes (which can be changed by the user) with a P-value less than 0.05 are selected. The specific functions of each component and the overall contribution of the generic approach are discussed in more detail in the following sections.

### The Grouping Component

The Grouping component (denoted as G and depicted as the orange section in Figure 4) is the first component of the G-S-M Tool. It is responsible for dividing the control variables into smaller groups based on pre-existing knowledge. In this case, the G component uses the grouping file provided by the user to create groups of attributes based on their known associations with specific diseases. Table X provides an example (GEO gds1962 dataset^17^) of these groups, including the name of the group name, the set of attributes associated with the group. The grouping component is a crucial step in the G-S-M Tool workflow because it helps to focus the analysis on specific sets of predictors that may be relevant to the groups of interest.

**Figure 4:**
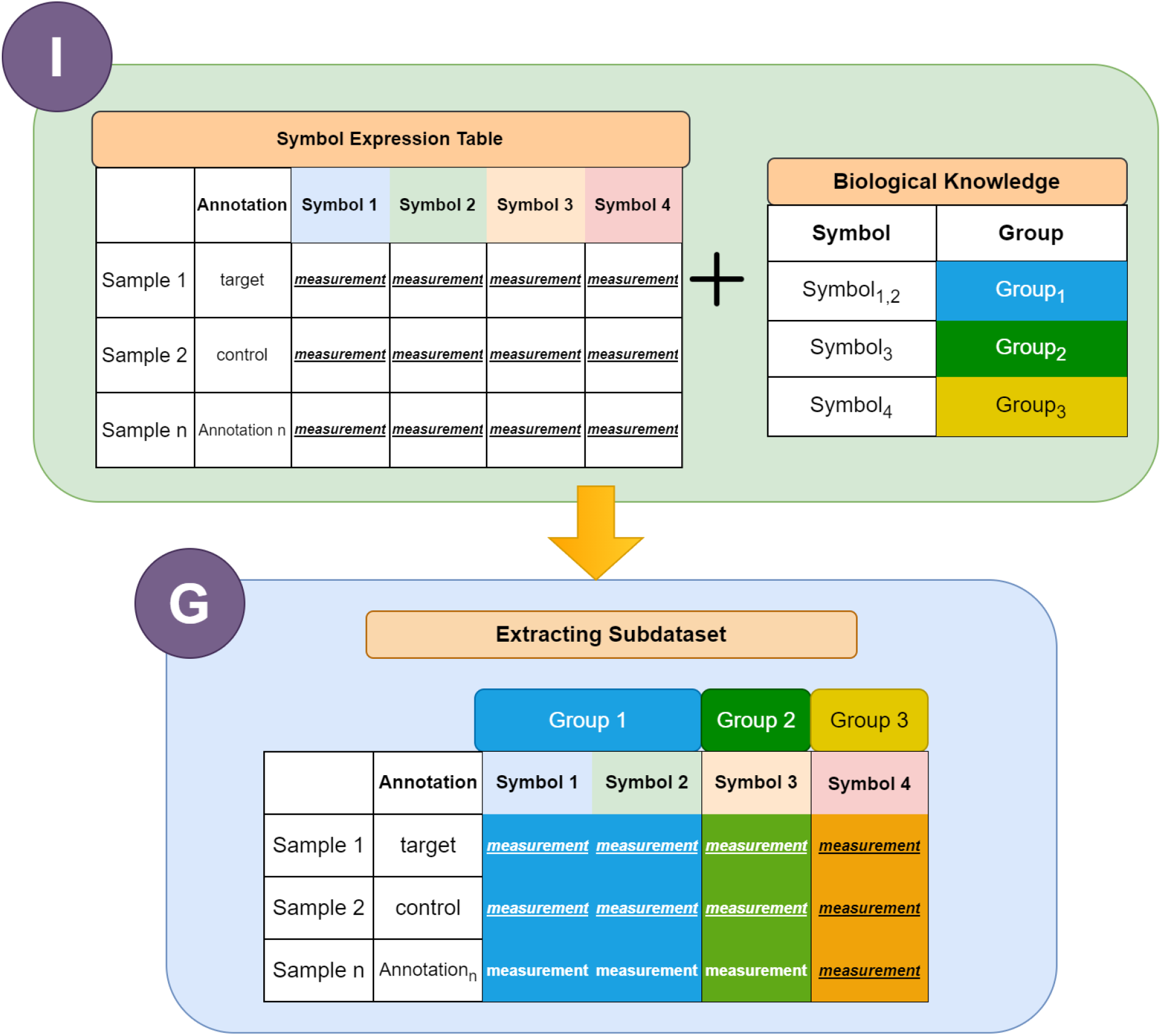
It represents the generation of subsets of the data based on specific disease groups. “I” refers to input and “G” refers to the Grouping module.

**Figure 5.**
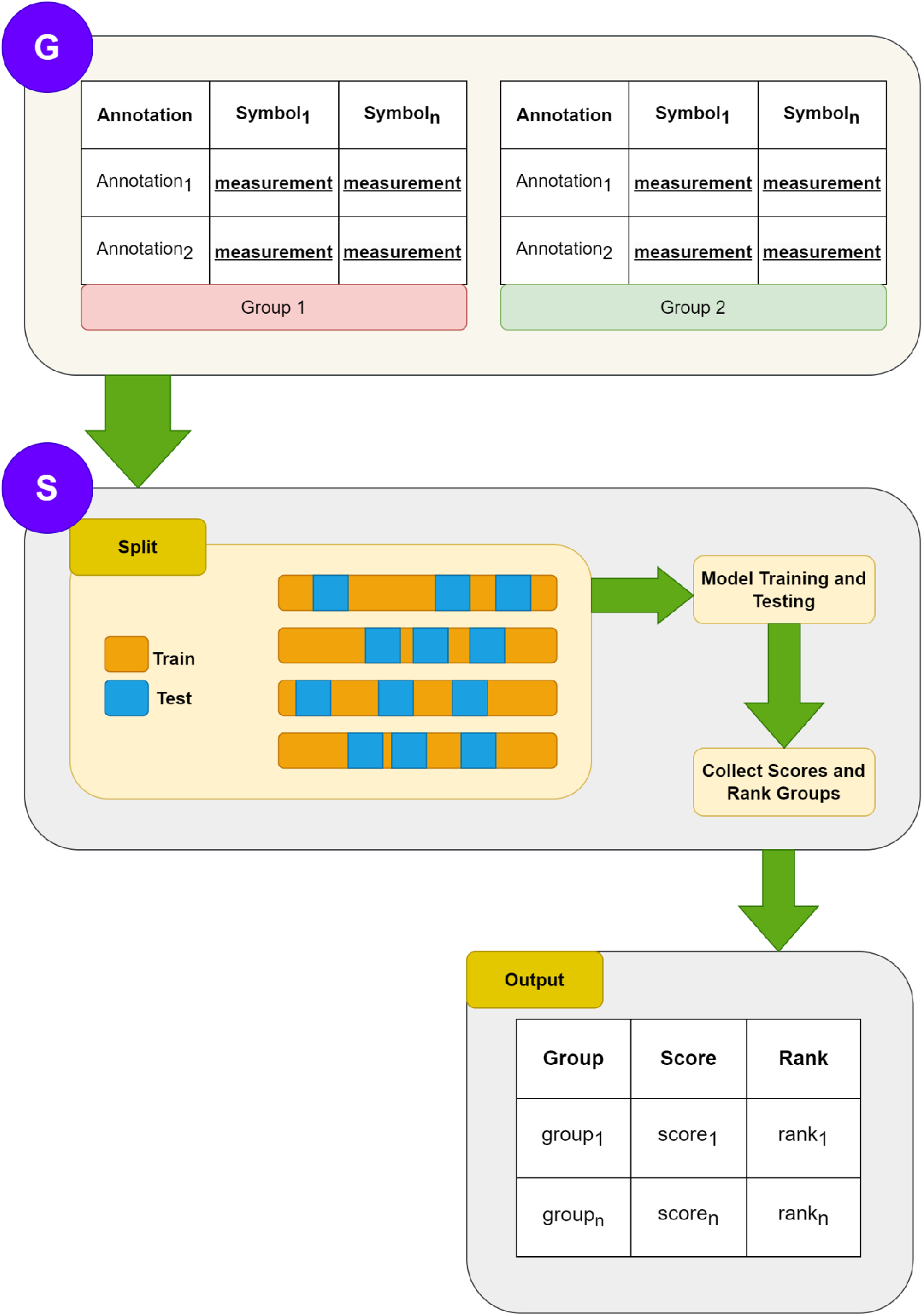
shows that The S component is applied to each of the sub-datasets within the G panel, which contains all the sub-datasets that have been divided into two groups based on disease.

### The scoring component

The G component generates m groups, which are subsequently scored by the S component based on their importance. To determine this score, the training data is randomly partitioned into a 90% train set and a 10% test set. The selected classifier is trained on the train set and used to make predictions on the test set, resulting in various statistics, including accuracy.

This process is repeated 5 times, and the average accuracy is taken as the final score for each group. The groups are then ranked in a table based on their scores, with the highest-scoring groups listed first. This table is then passed on to the M component.

### The Modeling Component

For the M component training, the top 10 groups from the S component were considered. In each iteration, the predictors from the top j groups were extracted from the data (e.g. when j=1, the group with the best result is used. When j=5, predictors in the top 5 groups are used.) and used as input features for the model, which could be one of several types (such as Decision Tree, Random Forest, Support Vector Machine (SVM), Naive Bayes, or Random Forest Based Treebag). The sub-data generated from this process was then evaluated using the test data and the resulting statistics were recorded in a table.

### Classifiers

A Decision Tree Classifier is a supervised learning algorithm that is used for classification tasks. It works by recursively partitioning the data into smaller subsets based on the feature values. The final partitions represent the different classes, and a tree structure is created to represent the decision process. Decision trees are easy to understand and interpret, and they can handle both numerical and categorical data. However, they can be sensitive to small variations in the data, which can lead to overfitting.

Support Vector Machine Classifier is a supervised algorithm that is used for classification and regression tasks. It works by finding the best boundary, called the hyperplane, that separates the different classes in the data. The data points closest to the hyperplane, called support vectors, have the greatest impact on the position of the hyperplane. SVM is effective in high-dimensional spaces and can handle non-linearly separable data through the use of kernel functions. However, it can be sensitive to the choice of kernel and the values of the parameters.

Naive Bayes Classifier is a probabilistic algorithm that is used for classification tasks. It works by using the Bayes theorem to estimate the probability of a class given the feature values. It assumes independence between the features, which is called the naive assumption. Naive Bayes is simple, fast, and requires small amounts of data. However, it can be affected by the presence of irrelevant features and the assumptions of independence.

Random Forest Classifier is an ensemble algorithm that is used for classification and regression tasks. It works by creating multiple decision trees and combining their predictions. Each tree is built using a random subset of the features, which helps to reduce the correlation between the trees. Random Forest is robust to overfitting, easy to interpret, and can handle high-dimensional data. However, it can be computationally expensive and require large amounts of memory.

Random Forest-based Treebag Classifier is an ensemble algorithm that is similar to Random Forest, but it uses Treebag Classifier. The Treebag Classifier is an algorithm that was developed to improve the performance of the Random Forest Classifier. It also uses multiple decision trees and combines their predictions, but it uses a bootstrap aggregation technique to improve the accuracy of the predictions. Like Random Forest, it is robust to overfitting, easy to interpret, and can handle high-dimensional data. However, it may also be computationally expensive and require large amounts of memory.

### Implementation of the G-S-M Tool

The G-S-M Tool tool has been developed using the KNIME platform, a widely utilized and highly versatile data analytics software known for its user-friendly graphical interface. The integration capabilities of KNIME, including the ability to incorporate scripts written in Python, R, and Java, have enabled the creation of a sophisticated and customizable workflow within the platform for implementing the G-S-M Tool algorithm. This workflow is composed of discrete nodes, each with designated functionalities, as well as meta-nodes, which are conglomerations of nodes that perform specific tasks as a unit.

The KNIME workflow for G-S-M Tool is a powerful data analysis tool that allows users to input their desired hyperparameters and then systematically process and analyze datasets. The workflow begins by utilizing the “List Files/Folders” node to upload a list of dataset names. It then iterates through each dataset, utilizing the “Table Reader” node to read the data and the “MissingValues” meta-node to filter out rows with missing values. The resulting filtered data is then input into the G-S-M meta-node for further processing and analysis. Overall, this workflow allows for efficient and comprehensive data analysis using the G-S-M Tool algorithm within the user-friendly and customizable environment provided by the KNIME platform.

### Pre-Start Requirements

#### KNIME Version

This workflow was made in version “4.6.3” of KNIME. Hence, utilizing a version that is at or above the specified version will serve to mitigate the occurrence of unnecessary errors.

#### Environment Settings

The G-S-M workflow incorporates both Python and R scripts. Therefore, to prevent any errors from occurring, it is essential to properly configure the KNIME Python settings by following the path:

File -> Preferences -> KNIME (left side of the pop-up) -> Python

Additionally, the R server must be running simultaneously during the execution of the workflow. To initiate the R server, the following commands should be executed in R/RStudio:

library(Rserve);

Rserve(args = “--vanilla”)

#### Knime Workflow

By adhering to these environmental settings, errors will be effectively mitigated in this stable version of the workflow.

- List Files/Folders Node (“Node 479”): needs to give the path of the directory which contains data
- Table Reader (“Node 478”): finds the “.table” file in a given directory and reads those values and put it into a table.
- Table Reader (“Read miRNA data”): finds the “.table” file in a given directory and reads those values and put it into a table.
- Row Filter (“remove 5.0”): If there is other than a value except for two classes one needs to remove that value from the class column.
- MissingValues (“Node 477”): extracts missing values and converts required columns from strings to numbers.
- User Inputs: component including Number of Iteration, Output folder name, Group File Path, and Model parameters.
- Set Up Parameters (“Node 433”): component including the number of initial clusters, the initial number of genes, the label of the positive class, the number of iterations for internal rank, and reduction ratio parameters.
- Rank Functions Weights (“Node 434”): Component including accuracy, sensitivity, specificity, F-measure, AUC, and precision weights.
- Upload Groups File (“Node 466”): uploading grouping file which comes from the “User Inputs” component.
- G-S-M (“Node 468”): that G-S-M operations are made.
- Display Result (“Node510”): Component that results are visualized.

#### Including Data Path

The source-choosing operation shown in Figure 6 should be performed to specify the input data to be used.”

**Figure 6.**
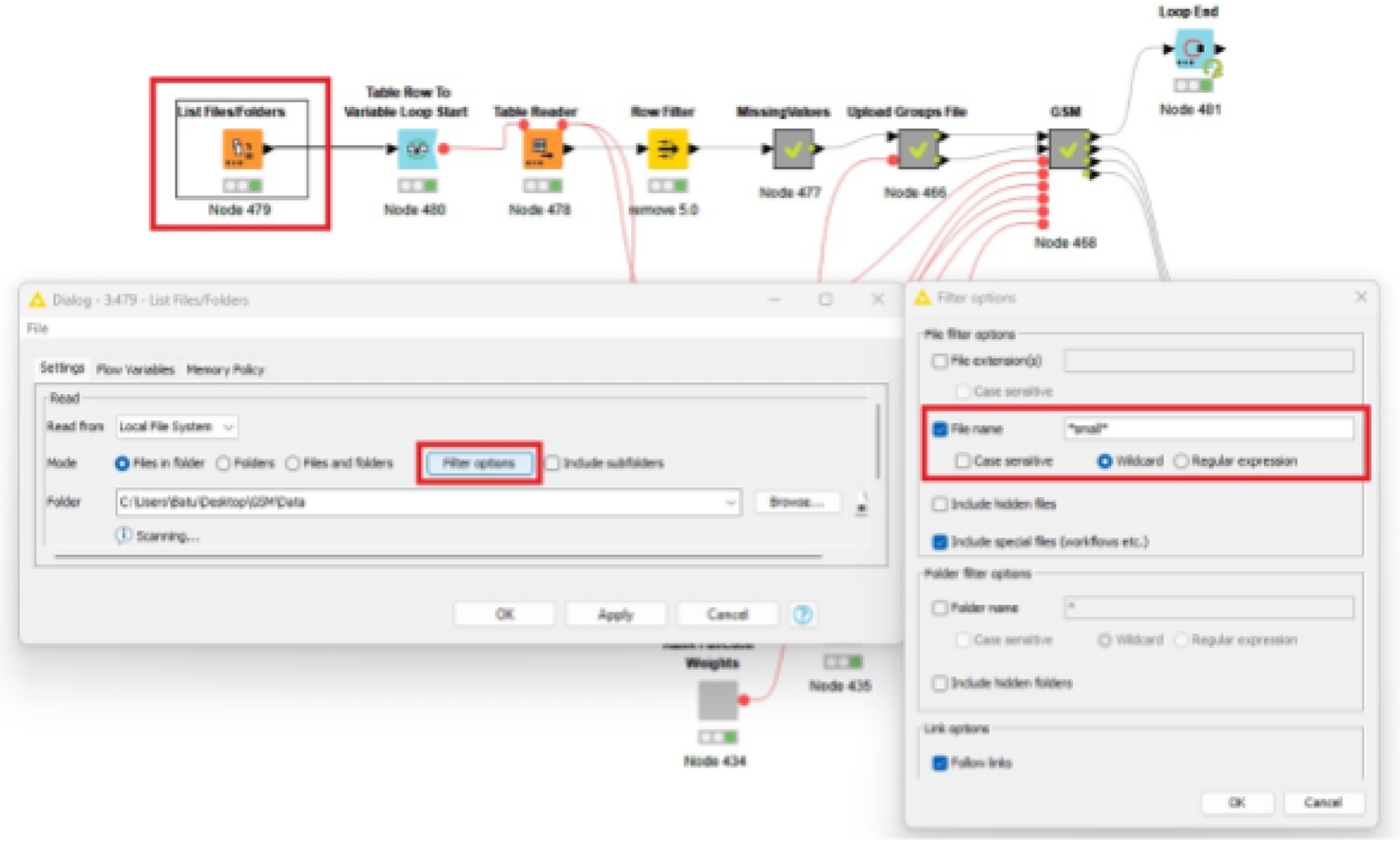
shows how a file can be specifically selected.

**Figure 7.**
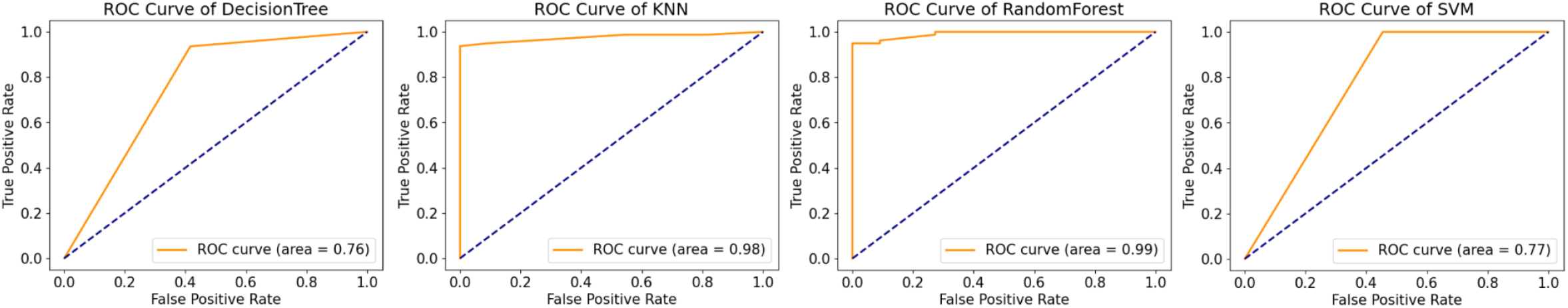
ROC results of the different models for GDS1962 dataset.

### Parameter Settings

The components below should be set before running the workflow.

#### User Inputs

This component possesses four distinct parameters.

1. Number of Iterations: determines how many iterations the model will make
2. Output Folder Name: sets the name of the output folder name
3. Choose Group File: requests file found in gene clusters
4. Choose Model Name: allows to specify the model to be selected by the user (Current models are: Random Forest, Support Vector Machine, Decision Tree, Naive Bayes, and Treebag Classifiers)

#### Set Up Parameters

This component contains the following 5 different parameters.

1. Number of Initial Clusters
2. Initial Number of Genes: Determines the number of genes used in the t-test.
3. The Label of the Positive Class: Requests the value name of positive targets.
4. Number of Iterations for Internal Rank: (Default value: 5)
5. Reduction Ratio: (Default value: 0.1)

#### Rank Function Weights

The parameters in this component are used to establish the relative importance of different factors in the formula used to calculate the ranking.

1. Accuracy Weight: (Default value: 1.0)
2. Sensitivity Weight: (Default value: 0.0)
3. Specificity Weight: (Default value: 0.0)
4. F-measure Weight: (Default value: 0.0)
5. AUC Weight: (Default value: 0.0)
6. Precision Weight: (Default value: 0.0)

## Results

### Reproduction of Previous Results

The generalized G-S-M approach should be able to reproduce the results from our previous works such as maTE, GenOntology, and PriPath.We selected three of our previous works to test whether the generalized implementation is able to reproduce the results. Clearly, there is an element of randomness in the MCCV part of the algorithm, so a perfect reproduction cannot be expected. However, we expect the same groups to occur in approximately the same sequence for datasets that were previously separable, depending on the difference in previous average score. However, we do not expect to reproduce the scores exactly.

### Model Performance Evaluation

Many parameters influence the performance of the G-S-M approach. The selection of the training algorithm also has a large effect. KNIME offers a wide range of ML algorithms and changing the algorithm in our workflow is easy. We provide a video for the purpose on our GitHub repository (https://www.youtube.com/watch?v=2Z5McuzIL7M). In the following, we assess the effect of different parameters and the selection of different ML algorithms on the G-S-M performance. We measure the performance by the average accuracy achieved by the best model.

When working with large datasets, it is common, in biology, to have a large number of features. However, not all features may be relevant for the specific task. Feature selection can be used to reduce the number of attributes, but these methods can be computationally expensive and time-consuming, especially when dealing with large datasets. To identify the most relevant features effectively, feature selection can be applied. This process involves evaluating each feature based on its ability to predict the target variable and ranking them accordingly. By selecting a subset of the top-ranking features, the most informative features can be selected while disregarding the less important ones. In this case, the user can select any number of features by applying feature scoring.

To address the issue of imbalanced classes in the target variable, under-sampling of the majority class was employed during the training phase. This technique aims to balance the distribution of the classes in the training data, thus reducing the potential for bias in the model.

To evaluate the performance of the model, a variety of quantitative metrics were utilized, including accuracy, specificity, sensitivity, and the area under the curve (AUC) of the receiver operating characteristic (ROC).

To further improve the robustness of the results, an n-fold (n is a user choice) Monte Carlo cross validation approach was employed. This technique involves randomly dividing the data into training and testing sets in each iteration, and then evaluating the model’s performance using the metrics mentioned above. By performing this process repeatedly, we gain a better understanding of how the model might generalize.

Our results from ROC analysis indicate that the Support Vector Machine (SVM) model had a ROC score of 1.0, and the Naive Bayes model also had a ROC score of 1.0, indicating that they performed well in classifying the binary-labeled target. The Random Forest model had a ROC score of 0.9, and The TreeBag model also had a ROC score of 0.9, indicating that they also performed well but not as well as the SVM and Naive Bayes models. However, the Decision Tree model had a ROC score of 0.75, indicating that it performed worse than the other models. This tool can be a useful resource for researchers and can help to improve the accuracy and effectiveness of their models.

In addition to evaluating the performance of the models, the study also aimed to generate lists of disease groups and corresponding genes that are relevant to the target variable. To do this, a prioritization approach was used to rank the lists generated by the model in each iteration. To account for the potential variability of the ranking across different iterations, rank aggregation techniques were employed. Specifically, the RobustRankAggreg R package11 was integrated into the G-S-M Tool workflow. The RobustRankAggreg algorithm assigns a p-value to each element in the aggregated list, which reflects the quality of the ranking of each element or entity compared to the expected value. This helps to provide a more comprehensive and robust understanding of the relationships between the target variable and the disease groups and genes identified by the model.

## Discussion and Conclusion

In the Discussion section, the results of the ROC analysis demonstrate the efficacy of the Support Vector Machine (SVM) and Naive Bayes models in classifying binary-labeled targets, as evidenced by their ROC scores of 1.0. These results highlight the potential of these models in tasks related to diagnosis, prognosis, treatment, and biological studies. The Random Forest and TreeBag models, with ROC scores of 0.9, also showed promising performance and could be considered for similar tasks. However, the Decision Tree model, with a ROC score of 0.75, exhibited a lower performance compared to the other models. These findings suggest that alternative models may be more suitable for tasks that require high accuracy in binary classification.

The G-S-M tool, which combines machine learning and prior knowledge to the group and score features, has been shown to effectively identify the top-performing feature groups. These feature groups can be used to train machine-learning models, thereby enhancing the accuracy and efficacy of models used in tasks in studies. Furthermore, the G-S-M tool enables a more comprehensive understanding of the underlying mechanisms of a given system through the integration of machine learning and prior knowledge, potentially leading to novel insights and discoveries.

In conclusion, the G-S-M tool presents a valuable resource for researchers in various fields. By effectively identifying relevant features, the G-S-M tool can improve the accuracy and effectiveness of models used in tasks. The results of the ROC analysis demonstrate that the SVM and Naive Bayes models are the most suitable for binary classification tasks in the dataset used, while the Random Forest and TreeBag models also showed promising performance. On the other hand, the Decision Tree model exhibited lower performance. The G-S-M workflow is made available for download, making it readily accessible to researchers in the field. This tool can be a powerful tool to improve the accuracy and effectiveness of models used in the studies.

## Data Availability

All datasets used in this study are publicly available in the GEO database (https://www.ncbi.nlm.nih.gov/geo/) via the ID provided with each of the datasets. The KNIME workflow and all test input files are available on our GitHub repository: https://github.com/malikyousef/The-G-S-M-Grouping-Scoring-Modeling-Approach.

